# Cardiac pericytes are necessary for coronary vasculature integrity and cardiomyocyte differentiation

**DOI:** 10.1101/2022.08.29.505703

**Authors:** Lauriane Cornuault, François-Xavier Hérion, Paul Rouault, Ninon Foussard, Philippe Alzieu, Candice Chapouly, Alain-Pierre Gadeau, Thierry Couffinhal, Marie-Ange Renault

## Abstract

**Introduction**: While the critical role of pericytes in maintaining vascular integrity has been extensively demonstrated in the brain and in the retina, very little is known about their role in the heart. **Objective**: We aim to investigate structural and functional consequences of partial pericyte depletion (about 60%) in the heart of adult mice. **Methods**: To deplete pericyte in adult mice, we used Pdgfrb-Cre/ERT2; Rosa-DTA mice and compared their phenotype to the one of control mice (Rosa-DTA) chosen among their littermates. Cardiac function was assessed via echocardiography and left ventricle (LV) catheterization one month after the first tamoxifen injection. **Results**: Mice depleted with pericytes displayed increased coronary endothelium leakage and activation which was associated with increased CD45 + cell infiltration in the heart. Pericyte depletion also modified the phenotype of cardiomyocytes with an increased expression of Myosin Heavy Chain 7, a decreased expression of ATPase Sarcoplasmic/Endoplasmic Reticulum Ca2+ Transporting 2, Connexin 43 and a decreased phosphorylation of Phospholamban suggesting cardiomyocyte dedifferentiation and impaired contractility. As a consequence, mice depleted with pericytes had a reduced LV ejection fraction and an increased end-diastolic pressure demonstrating both systolic and diastolic dysfunction. Accordingly, mice depleted with pericytes presented a decreased LV contractility and an increased LV relaxation time (dP/dt_min_). Besides this study reveals that cardiac pericytes may undergo strong remodeling upon injury. **Conclusion**: Cardiac pericyte depletion induces both systolic and diastolic dysfunction suggesting that pericyte dysfunction may contribute to the occurrence of cardiac diseases.

## Introduction

Pericytes are perivascular cells that reside in the microvasculature and share a basement membrane with the underlying endothelial cells. Pericyte density varies between different organs, in the brain, the endothelial cell–pericyte ratio varies between 1:1 and 3:1. It represents the organ with the highest density of pericytes (Armulik, Abramsson, and Betsholtz 2005). Accordingly, the brain is also the organ in which the role of pericyte has been the most investigated. Notably, in this organ, pericytes are thought to play a role in regulating permeability across the blood–brain barrier (Armulik et al. 2010; Daneman et al. 2010) since mice deficient in pericytes or exhibiting decreased pericyte function have increased vascular permeability (Armulik et al. 2010; Daneman et al. 2010). Notably, pericyte function and maintenance depends on PDFGB expression by quiescent adult microvascular brain endothelium (Vazquez-Liebanas et al. 2022).

The pericyte content of the cardiac micro-vasculature is thought to be closer to the one of the cerebral vasculature with an endothelial–pericyte ratios of 2:1–3:1 (Su et al. 2021). It is also likely that pericytes contribute to the maintenance of microvascular function in the heart given that loss of pericytes has been associated with increased vascular permeability (Chintalgattu et al. 2013) (Siao et al. 2012), endothelial activation (Siao et al. 2012) and altered vasomotricity (Chintalgattu et al. 2013). Notably in mice treated with PDGF receptor inhibitors (sunitinib and CP-673451), decreased pericyte coverage of cardiac capillaries was associated with increased permeability, impaired endothelium-dependent vasodilation and decreased coronary flow reserve (Chintalgattu et al. 2013). Besides, expression of a mutant form of ProNGF, unable to be matured and which may target pericytes, was shown to induce endothelial activation and vascular leakage (Siao et al. 2012). Finally, pericyte loss observed in both *Sirt3* (He, Zeng, and Chen 2016) and *Notch3* (Tao et al. 2017) deficient mice was associated with decreased coronary flow reserve. Importantly, in each of these studies impaired microvascular integrity and function has been also associated with cardiac dysfunction, Notably, administration of the PDGF receptor specific inhibitor CP-673451 was shown to decrease ejection fraction (EF) (Chintalgattu et al. 2013), expression of the mutant form of ProNGF induced dilated cardiomyopathy, cardiac fibrosis, and contractile dysfunction (Siao et al. 2012), *Sirt3* deficient mice showed decreased EF and fractional shortening after myocardial infarction (He, Zeng, and Chen 2016) while *Notch3* deficient mice were shown to exhibit cardiac hypertrophy (Tao et al. 2017).

Altogether these studies suggest that pericyte may be beneficial to the coronary vasculature which may have an impact directly or indirectly on cardiac function. However, in the heart pericytes were also proposed to have deleterious effects. Notably, pericytes may become myofibroblasts and contribute to fibrosis. (Greenhalgh, Iredale, and Henderson 2013). Also, capillary pericytes are suggested to be responsible for coronary no-reflow after myocardial ischemia by inducing capillary constriction (O’Farrell et al. 2017) via a mechanism involving GPR39 (Methner et al. 2021, 39).

In pathophysiological conditions, cardiac pericytes have been either shown to be decreased or increased in number. Loss of pericytes has been shown to constitute a remarkable feature of aging in several organs including the heart in both mice and humans (Chen et al. 2021). Also High-fat diet was shown to induce pericyte loss in the heart of mice (Zeng et al. 2015). On the contrary, cardiac pericytes were shown to be disorganized and more numerous in obese and hypertensive ZSF1 rats (van Dijk et al. 2016). In other conditions, such as SARS-CoV-2 infection, the phenotype of pericytes is proposed to be modified by the acquisition of a more contractile phenotype (Robinson et al. 2020). Finally, pericyte dedifferentiation leading to increased endothelial permeability is proposed to be induced by the downregulation of Hypoxia-Induced Endoplasmic Reticulum Stress Regulating lncRNA in human heart failure (Bischoff et al. 2017).

In conclusion, cardiac pericytes may participate to the pathophysiology of cardiac diseases through a wide range of mechanisms. However, studies performed so far are mainly descriptive and conclusions made are based on associations. The purpose of the present study is to investigate the specific consequences of pericyte depletion on cardiac structure and function.

## Methods

### Mice

B6.Cg-Tg(Pdgfrb-cre/ERT2)6096Rha/J (strain# 029684), Gt(ROSA)26Sortm4(ACTB-tdTomato,- EGFP)Luo/J (Strain # 007576) and B6.129P2-Gt(ROSA)26Sortm1(DTA)Lky/J (Strain#009669) were obtained from the Jackson laboratory.

Animal experiments were performed in accordance with the guidelines from Directive 2010/63/EU of the European Parliament on the protection of animals used for scientific purposes and approved by the local Animal Care and Use Committee of Bordeaux University. Both males and females were used. Mice were either sacrificed by cervical dislocation or exsanguination under deep anesthesia (ketamine 100 mg/kg and xylazine 20 mg/kg, IP).

### Echocardiography

Left-ventricular ejection fraction and LV dimension were measured on a high-resolution echocardiographic system equipped with a 30-MHz mechanical transducer (VEVO 2100, VisualSonics Inc.) as previously described (Renault et al. 2009; Roncalli et al. 2011). Mice were anchored to a warming platform in a supine position, limbs were taped to the echocardiograph electrodes, and chests were shaved and cleaned with a chemical hair remover to minimize ultrasound attenuation. UNI’GEL ECG (Asept Inmed), from which all air bubbles had been expelled, was applied to the thorax to optimize the visibility of the cardiac chambers. Ejection fractions were evaluated by planimetry as recommended (Schiller et al. 1989). Two-dimensional, parasternal long-axis and short-axis views were acquired, and the endocardial area of each frame was calculated by tracing the endocardial limits in the long-axis view, then the minimal and maximal areas were used to determine the left-ventricular end-systolic (ESV) and end-diastolic (EDV) volumes, respectively. The system software uses a formula based on a cylindrical-hemiellipsoid model (volume=8.area^2^/3π/ length) (Mohan et al. 1992). The left-ventricular ejection fraction was derived from the following formula: (EDV- ESV)/EDV*100. The cardiac wall thickness, left ventricular posterior wall (LVPW), inter-ventricular septum (IVS) and left ventricular internal diameter (LVID) were calculated by tracing wall limits in both the long and short axis views.

### LV pressure /systolic blood pressure measurement

LV diastolic pressure measurement was assessed via invasive catheterization technique. Briefly, mice were anesthetized with Isoflurane. A Scisense pressure catheter (Transonic) was inserted into the LV through the common carotid artery. Pressure was recorded using LabChart software. End diastolic pressure, dP/dt minimum and maximum, Tau and heart rate were automatically calculated by a curve fit through end-systolic and end-diastolic points on the pressure plot.

### Blood sampling for biochemical marker analysis/NFS

Blood samples were collected by the heparin retroorbital bleeding method at sacrifice. Blood cell counts were determined using an automated counter (scil Vet abc Plus+). Plasma was separated by a 10-min centrifugation at 2500 g and then stored at -80°C. Concentrations of the following biomarkers were measured using an Architect CI8200 analyzer (Abbott Diagnostics, North Chicago, Illinois, USA): triglycerides, using the lipoprotein-lipase/glycerol kinase/oxidase/peroxidase method; total cholesterol, using the esterase/oxidase/peroxidase method; and HDL cholesterol, using the accelerator/selective detergent/esterase/oxidase/peroxidase method. LDL cholesterol was then estimated using the Friedewald formula (LDL cholesterol [mmol/L] = total cholesterol – HDL cholesterol – [triglycerides/2,2], or LDL cholesterol [mg/dL] = total cholesterol – HDL cholesterol – [triglycerides/5]).

### Tissue staining/Immunostaining

Capillary density was evaluated in sections stained with anti-CD31 or anti-Podocalyxin (PODXL) antibodies by quantifying the number of CD31+ or PODXL+ elements. Muscularized vessels were identified using anti–alpha-smooth muscle actin (SMA) antibodies. Pericyte were identified using anti-NG2 or anti-PDGFRB antibodies. Leucocyte, macrophage, T-cell and B-cell infiltrations were quantified using anti-CD45, anti-CD68, anti-CD3 and anti-B220 antibodies, respectively. Endothelial adherens junction was characterized using anti-CDH5 antibodies. Edema was measured after FGB staining of muscle sections and quantified as the mean GFB-positive areas. (See Supplemental table 1 for antibody references.)

Cardiomyocyte mean surface area was measured using ImageJ software after membrane staining with Wheat Germ Agglutinin (WGA), Alexa Fluor™ 488 Conjugate (Invitrogen). Fibrosis was assessed after sirius red staining of heart sections by quantifying the percentage of red stained area.

Quantifications were done on images were acquired using an Axiozoom V16 or axioscope A1 (Zeiss) under 200x magnification. They were conducted on 10 randomly selected images were performed using ImageJ/Fiji v2.0.0-rc-59 software (National Institute of Health, USA) by an investigator blinded to genotype. More precisely, each sample received a unique number. At the end of the experiment, the genotype/treatment for each sample was unveiled to enable data analysis.

For immunohistochemical analyses, primary antibodies were sequentially coupled with biotin-conjugated secondary antibodies and streptavidin-horseradish peroxidase (HRP) complex (Amersham). Staining was then revealed with a DAB substrate kit (Vector Laboratories) and tissue sections were counterstained with hematoxylin (see Supplemental table 2 for secondary antibody references).

For immunofluorescence analyzes, primary antibodies were resolved with Alexa-Fluor–conjugated secondary polyclonal antibodies (see Supplemental table 2). Nuclei were counterstained with DAPI (4′,6-diamidino-2-phenylindole).

For both immunohistochemical and immunofluorescence analyses, negative control experiments to check for antibody specificity were performed using secondary antibodies.

### Cell culture

Human cardiac microvascular endothelial cells (HMVEC-C) (Lonza) were cultured in endothelial basal medium-2 (EBM-2) supplemented with EGM™-2 BulletKits™ (Lonza). Cells at passage 3 were used.

Mouse pericytes were isolated from 10 day-old pups. Briefly, hearts were dissociated using 2 mg/mL type IV collagenase (Gibco™, ThermoFisher) for 1 hour at 37°C and the resulting dissociated cells were filtrated on a 30 µm strainer. Pericytes were labelled with rat anti-mouse CD146 microbeads (Miltenyi Biotec). Labelled cells were then isolated magnetically, plated and cultured in pericyte Growth Medium 2 (PromoCell). Cells at passage 1 and 2 were used.

HMVEC-C and mouse pericytes were co-cultured in 0.4 µm-pored Transwells in 6-well plates (Corning)

### Quantitative RT-PCR

RNA was isolated using Tri Reagent® (Molecular Research Center Inc) as instructed by the manufacturer, from heart tissue that had been snap-frozen in liquid nitrogen and homogenized. For quantitative RT-PCR analyses, total RNA was reverse transcribed with M-MLV reverse transcriptase (Promega) and amplification was performed on an AriaMx Real Time PCR system (Agilent Technologies) using B-R SYBER® Green SuperMix (Quanta Biosciences). Primer sequences are reported in Supplementary table 3.

The relative expression of each mRNA was calculated by the comparative threshold cycle method and normalized to 18S rRNA expression.

### Western blot analysis

ICAM-1 and VCAM-1 protein levels were evaluated by SDS PAGE using goat anti-Icam-1 (R&D systems Cat# AF-796) and rabbit anti-Vcam-1 (abcam, Cat# ab134047) antibodies respectively. ATP2A2 protein level was evaluated by SDS PAGE using rabbit anti-ATP2A2 antibodies (Badrilla, Cat# A010- 80). PLN phosphorylation at Serine 16 and Threonin 17 was evaluated by SDS PAGE using rabbit anti-phospho-PLN Ser16 (Badrilla, Cat# A010-12), rabbit anti-phospho-PLN Thr17 (Badrilla, Cat# A010-13) and mouse anti-total PLN (Badrilla, Cat# A010-14).RYR2 phosphorylation was evaluated by SDS PAGE using rabbit anti-phosphoRYR2 antibodies (Badrilla, Cat# A010-31).

Human CDH5 and ICAM-1 protein levels were evaluated by SDS PAGE using rabbit anti-CDH5 (Cell signaling technology Cat# 2500) and mouse anti-ICAM-1 (Santa Cruz biotechnology, Cat# sc-8439) antibodies respectively.

Protein loading quantity was controlled using mouse monoclonal anti-α-tubulin antibodies (Sigma, Cat# T5168).

### Statistics

Results are reported as mean ± SEM. Comparisons between groups were analyzed for significance with the non-parametric Mann-Whitney test using GraphPad Prism v8.0.2 (GraphPad Inc, San Diego, Calif). The normality and variance were not tested. Differences between groups were considered significant when p≤0.05 (*: p≤0.05; **: p≤0.01; ***: p≤0.001).

## Results

### Cardiac capillaries are covered by SMA-, NG2+ pericytes

In the heart less than 2% of vessels, essentially arterioles, are covered by smooth muscle myosin heavy chain (smMHC) positive smooth muscle cells. Indeed, the vast majorities of capillaries, i.e. 95% of them, are covered by NG2 positive, SMA negative pericytes (Figure 1A-C). Cardiac pericytes are stellate shaped cells with cytoplasmic processes contacting several capillaries (Figure 1D-F).

**Figure 1.**
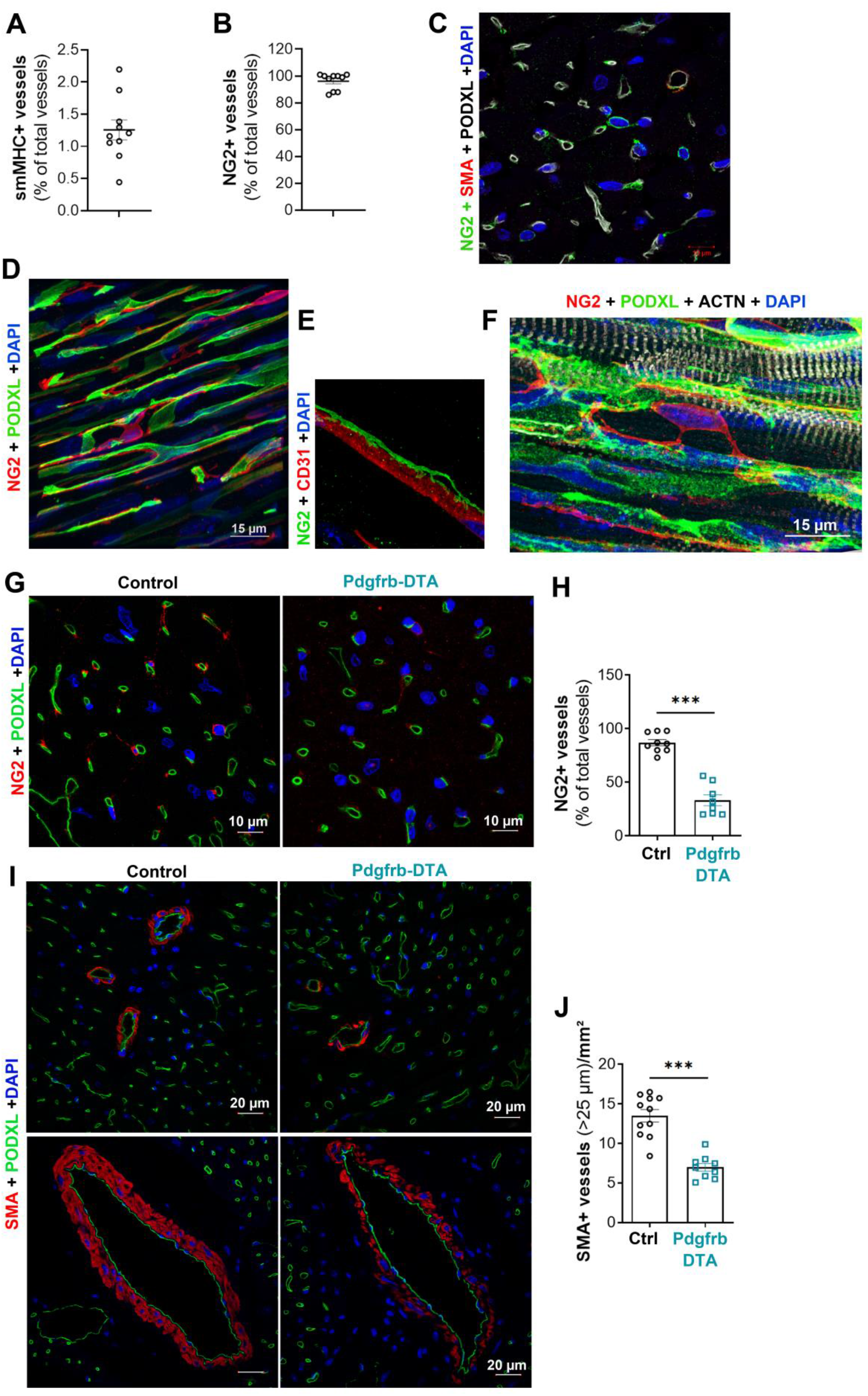
95 % of the cardiac vasculature is covered by pericytes. (A) Heart cross sections were co- immunostained with anti-smMHC and anti-PODXL antibodies to identify SMCs and ECs respectively. The percentage of smMHC+ vessels over total PODXL+ vessels was calculated. **(B)** Heart cross sections were co-immunostained with anti-NG2 and anti-PODXL antibodies to identify pericytes and ECs respectively. The percentage of NG2+ vessels over total PODXL+ vessels was calculated. (**C**) Heart cross section were co-immunostained with anti-SMA (in red), anti-NG2 (in green) and anti-PODXL (in white) antibodies to identify SMCs, pericytes and ECs respectively. (**D**) Heart thick sections were co- immunostained with anti-NG2 (in red) and anti-PODXL (in green) antibodies to identify pericytes and ECs respectively. (**E**) Heart thick sections were co-immunostained with anti-NG2 (in green) and anti- CD31 (in red) antibodies to identify pericytes and ECs respectively. (**E**) Heart thick sections were co- immunostained with anti-NG2 (in red), anti-PODXL (in green) and anti-ACTN2 (in white) antibodies to identify pericytes, ECs and cardiomyocyte respectively. (**G-J**) Pdgfrb-Cre/ERT2; RosaDTA (Pdgfrb-DTA) and RosaDTA (Control) mice were administered with tamoxifen. Mice were sacrificed 28 days after the first injection. (**G**) Heart cross sections were co-immunostained with anti-NG2 (in red) and anti- PODXL (in green) antibodies to identify pericytes and ECs respectively. (**H**) The percentage of NG2+ vessels over total PODXL+ vessels was calculated (n=8 and 9). (**I**) Heart cross sections were co- immunostained with anti-SMA (in red) and anti-PODXL (in green) antibodies to identify SMCs and ECs respectively. (**J**) The number of SMA+ vessels per mm^2^ was counted (n=9 and 11). ***: p≤0.001 (Mann Whitney test)

### Pdgfrb-DTA mice display a 60% pericyte depletion in the heart

To investigate the role of cardiac pericytes, we have chosen to measure the pathophysiological consequences of partial pericyte depletion on cardiac microvasculature, cardiomyocytes and heart function. To do so, we specifically induced diphtheria toxin (DTA) expression under the Pdgfrb promoter in adult mice. At first, we verified that Pdgfrb promoter was active in cardiac pericytes in adult mice, by crossing Pdgfrb-Cre/ERT2 mice with Rosa-mTmG mice. As shown in supplemental Figure 1A-B, GFP expression, in the heart was detected in most NG2+ pericytes but also, as expected, by SMA+ smooth muscle cells.

In the meantime, we have examined GFP expression in several other organs, notably the aorta, in which GFP expression was detected in few smooth muscle cells of the media, in the brain, in which GFP was detected in perivascular cells but also unexpectedly in glial cells, in the kidney in which GFP expression was detected in some SMA+ cells of arterioles but not in NG2 + cells within the glomerulus. Also GFP expression was detected in capillaries within the lacteals but not in SMCs surrounding the intestine. In the lung GFP expression was detected in both arterial and capillary mural cells, however, mural cells covering lung capillaries did not express NG2. Finally in the skeletal muscle, similarly to what we found in the heart, GFP expression was detected in every pericytes and few SMCs (Supplemental Figure 2).

Pericyte depletion was then induced by tamoxifen injections in 8 week-old Pdgfrb-Cre/ERT2; Rosa- DTA mice. Notably, to maintain constant pericyte depletion in the heart over time, we had to repeat tamoxifen administrations every 2 weeks. Indeed, in mice administered with tamoxifen once during 5 consecutive days, while the percentage of NG2 positive vessels was about 20% 7 days after the first tamoxifen injection, it was back to 90% after 28 days (Supplemental Figure 3A-B).

After 2 series of tamoxifen injections, i.e. one month after the first one, the percentage of cardiac capillaries covered by pericytes was reduced by about 60% in Cre positive mice compared to Cre negative mice (Figure 1G-H) and the number of arterioles covered by SMCs reduced by about 50% (Figure 1I-J) SMC coverage of coronary arteries was sparser (Figure 1I).

In parallel, we have examined the effectiveness of mural cell depletion in other organs. The most severe phenotypes were observed in the aorta, in which the media was strongly decellularized and in in the intestine lacteals and the skeletal muscle in which NG2+ pericytes have been totally depleted (Supplemental Figure 4). These “side” depletions induced significant weight loss (Supplemental Figure 5A-B) associated with circulating markers of malnutrition including decreased hemoglobin levels, triglyceride, cholesterol and proteins levels together with decreased Aspartate aminotransferase and Alkaline phosphatase levels (Supplemental Figure 5C-K). As expected, the smooth muscle cell loss in the aorta decreased diastolic blood pressure. Systolic blood pressure and heart rate were not modified (Supplemental Figure 5L-O). Finally, consistent with pericyte loss in the skeletal muscle, mice were severely intolerant to exercise (Supplemental Figure 5P).

Altogether these results demonstrate that Pdgfrb-DTA mice display successful pericyte depletion in the heart. However, considering the absence of pericyte specific marker and/or the absence of cardiac mural cell specific markers allowing pericyte depletion in the heart only, the interpretation of the data presented may have some limitations.

### Cardiac Pericyte are renewed

While performing a thorough histological analysis of the heart of Pdgfrb-DTA mice, we were surprised to see that the number of small vessels (with a diameter <25 µm) covered by SMA+ cells was significantly increased in the heart of Pdgfrb-DTA mice compared to control mice (Figure 2A-B). These SMA+ cells were actually NG2 positive, suggesting that the remaining pericytes in these mice display an activated phenotype characterized by SMA expression (Figure 2C). Besides, the heart of Pdgfrb-DTA mice displayed an increased number of Vimentin (VIM) positive mesemchymal cells (Figure 2D-E). Notably, among VIM positive cells, the number of PDGFRB negative cells (fibroblasts) did not increase (Figure 2F-G) while the number of PDGFRB positive cells significantly increased (Figure 2H). These results suggest that the number of cardiac fibroblasts does not change while the remaining pericytes became activated pericytes and proliferated. These PDGFB responsive cells did not all express NG2 (Figure 2H).

**Figure 2.**
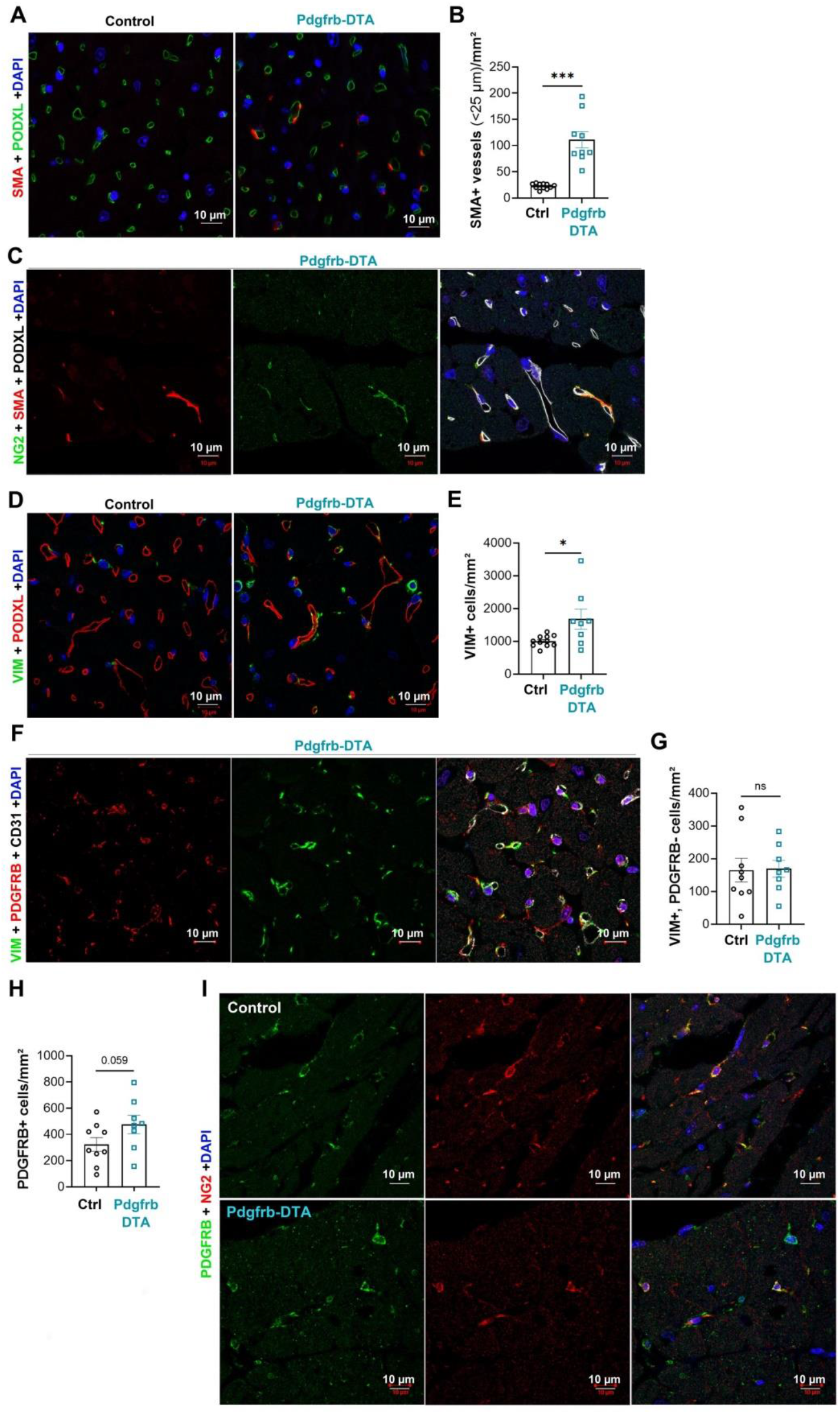
Pdgfrb-Cre/ERT2; RosaDTA (Pdgfrb-DTA) and RosaDTA (Control) mice were administered with tamoxifen. Mice were sacrificed 28 days after the first injection. (**A**) Heart cross sections were co-immunostained with anti-SMA (in red) and anti-PODXL (in green) antibodies to identify SMCs and ECs respectively. (**B**) The number of SMA+ vessels with a diameter ≤25 µm per mm^2^ was counted (n=9 and 11). (**C**) Heart cross sections were co-immunostained with anti-SMA (in red), anti-NG2 (in green) and anti-PODXL (in white) antibodies to identify SMCs, pericytes and ECs respectively. (**D**) Heart cross sections were co-immunostained with anti-VIM (in green) and anti-PODXL (in red) antibodies to identify fibroblasts and ECs respectively. (**E**) The number of VIM+ cells per mm^2^ was counted (n=9 and 11). (**F**) Heart cross sections were co-immunostained with anti-PDGFRB (in red), anti-VIM (in green) and anti-CD31 (in white) antibodies. (**G**) The number of VIM+, PDGFRB- cells per mm^2^ was counted (n=9 and 8). (**H**) The number of PDGFRB+ cells per mm^2^ was counted (n=9 and 8). (**I**) Heart cross sections were co-immunostained with anti-PDGFRB (in green), anti-NG2 (in red) antibodies. *: p≤0.05, ***: p≤0.001, ns: not significant (Mann Whitney test).

Accordingly, when we quantified cardiac fibrosis after Sirius red staining of cardiac sections we found that interstitial fibrosis was not significantly modulated; only perivascular fibrosis was (Supplemental Figure 6A-C). Moreover, the expression of several markers of fibrosis including Col1a1, Col3a1, Tgfb1 and Ctgf mRNA was not significantly different between Pdgfrb-DTA mice and control mice (Supplemental Figure 6D-G).

Altogether these results suggest that partial pericyte depletion induces activation and proliferation of the remaining pericytes.

### Pericyte depletion modifies the phenotype of cardiac capillary

Then we measured the consequences of mural cell depletion on cardiac microvasculature. As mentioned above, this was done one month after induction of pericyte depletion. As shown in figure 3A-B, pericyte depletion did not induce microvessel rarefaction, since capillary density was equivalent in both Pdgfrb-DTA and Ctrl mice. However, pericyte depletion led to an increased endothelial activation characterized by a significant increase in ICAM-1 and VCAM-1 expression. This phenotype was proved via immunostaining (Figure 3C-E) and confirmed by western blot analyses of total heart extracts (Figure 3F-H). Also pericyte depletion led to altered endothelial intercellular junctions since we observed that CDH5 staining was discontinuous in pericytes depleted mice (Figure 3I-J). We confirmed that endothelial intercellular junction integrity was altered in Pdgfrb-DTA mice since Fibrinogen (FGB) extravasation was significantly increased in Pdgfrb-DTA mice compared to control mice (Figure 3K-L) revealing that capillary permeability in abnormally increased in mice depleted with pericytes.

**Figure 3.**
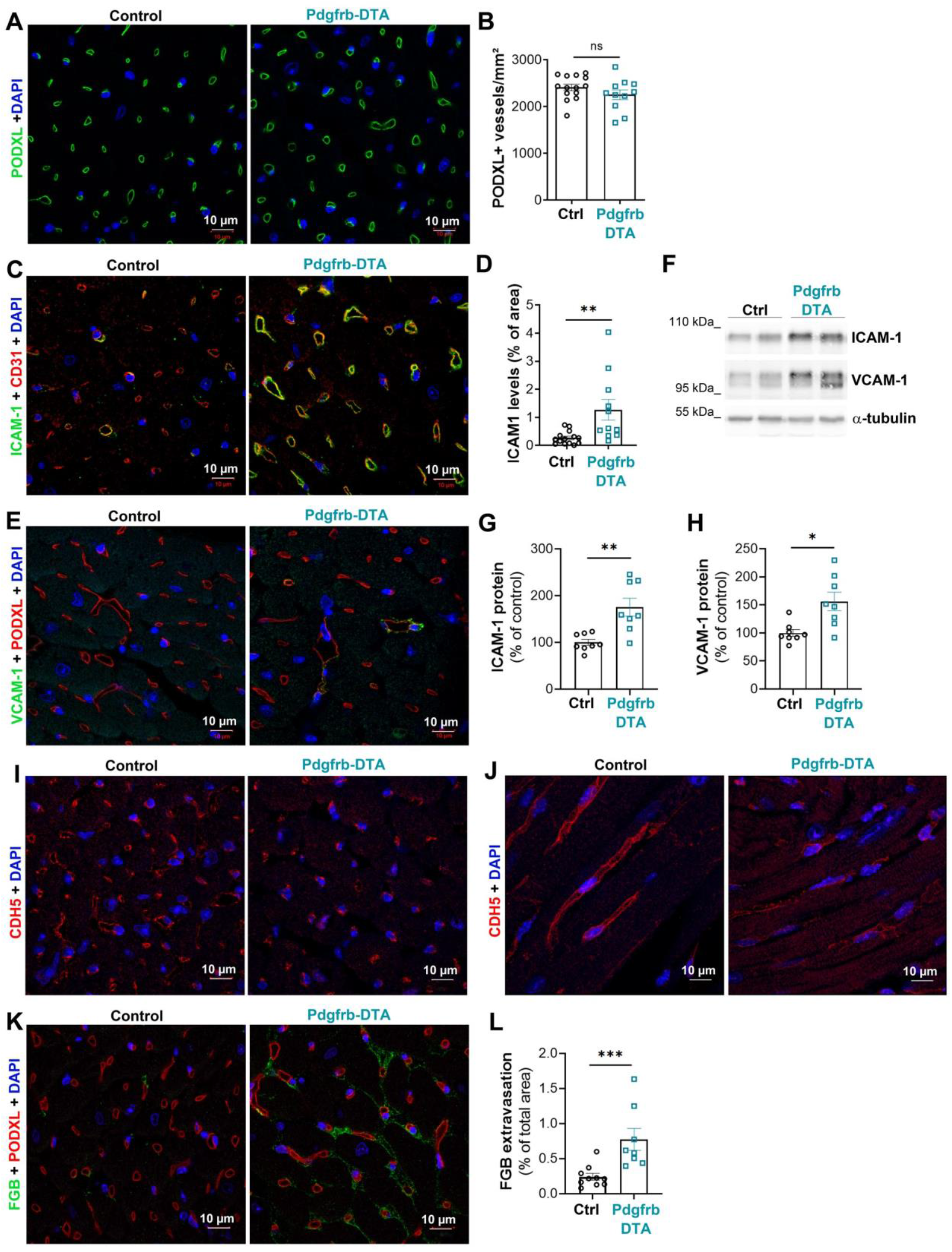
Pericyte are necessary to maintain cardiac capillary integrity. Pdgfrb-Cre/ERT2; RosaDTA (Pdgfrb-DTA) and RosaDTA (Control) mice were administered with tamoxifen. Mice were sacrificed 28 days after the first injection. (**A**) Heart cross sections were immunostained with anti-PODXL antibodies to identify ECs. (**B**) The number of PODXL+ capillary per mm^2^ was counted (n=14 and 11). (**C**) Heart cross sections were co-immunostained with anti-ICAM-1 (in green) and anti-CD31 (in red) antibodies. (**D**) ICAM-1 + surface area was quantified using Image J software (n=13 and 10). (**E**) Heart cross sections were co-immunostained with anti-VCAM-1 (in green) and anti-PODXL (in red) antibodies. (**F**) ICAM-1 and VCAM-1 protein expression was quantified in total heart extract by western blot analysis. ICAM-1 (**G**) and VCAM-1 (**H**) protein was quantified using image J software and normalized to α-tubulin (n=8 per group). (**I-J**) Heart cross sections were immunostained with anti- CDH5 antibodies to identify adherens junction. (**K**) Heart cross sections were co-immunostained with anti-FGB and anti-PODXL antibodies. (**L**) FGB+ surface area was measured using Image J software (n=10 and 8). *: p≤0.05, **: p≤0.01, ***: p≤0.001, ns: not significant (Mann Whitney test).

We confirmed the critical role of cardiac pericyte in maintaining endothelial integrity and immune quiescence in vitro. To do so we co-cultured mouse cardiac pericytes with human micro-vascular cardiac endothelial cells (HMVEC-C) (Supplemental figure 7A). HMVECs co-cultured with pericytes expressed significantly higher CDH5 protein level, while CDH2 mRNA was downregulated (Supplemental Figure 7B-D) signing increased endothelial differentiation. Consistent with in vivo data, pericytes prevented HMVEC-C activation by decreasing ICAM-1, VCAM-1 and IL-6 levels (Supplemental Figure 7E-I).

Consistent with endothelial activation Pdgfrb-DTA mice display cardiac inflammation with significantly increased Il-6 mRNA level (Supplemental Figure 8A) and increased CD45+ leucocyte infiltration (Supplemental Figure 8B-C) in the heart. Notably, neither CD68+ macrophage nor CD3+ T-cell infiltration was increased (Supplemental Figure 8D-H), while B220 B-cells infiltration was significantly increased (Supplemental Figure 8I-J).

Altogether these results confirm association studies and demonstrate that just like in the brain pericyte are necessary for cardiac capillary integrity and immune quiescence.

### Pericyte depletion modifies the phenotype of cardiomyocytes

To assess the phenotype of cardiomyocytes, we first performed WGA staining of heart cross sections and found that the cardiomyocytes did not show hypertrophy (Figure 4A-B). Consistent with this, the heart weight over tibia length ratio was not modified and so were cardiac dimensions (i.e. interventricular septum (IVS) and left ventricular posterior wall (LVPW) thickness and LV internal diameter (LVID) in diastole) measured via echocardiography (Figure 4D-F). However, WGA staining revealed that Pdgfrb-DTA had expanded cardiac interstitial space (Figure 4A), which is consistent with increased vascular leakage.

**Figure 4.**
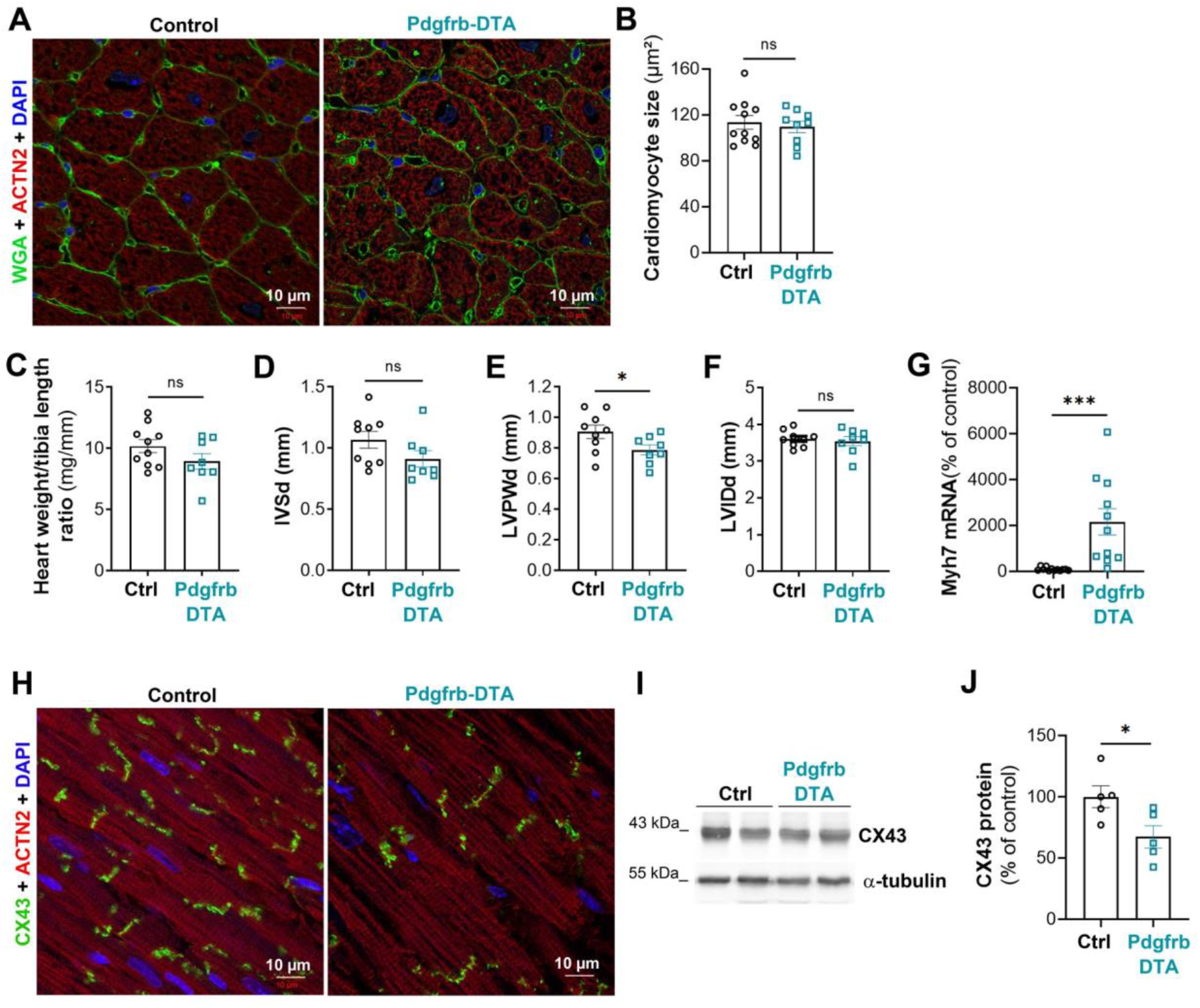
Pericyte depletion induces cardiomyocyte dedifferentiation but not cardiomyocyte hypertrophy. Pdgfrb-Cre/ERT2; RosaDTA (Pdgfrb-DTA) and RosaDTA (Control) mice were administered with tamoxifen. Mice were sacrificed 28 days after the first injection. (**A**) Heart cross sections were co-stained with WGA (in green) and anti-ACTN2 (in red) antibodies. (**B**) Cardiomyocyte cross section area was measured using Image J software (n=11 and 9). (**C**) The heart weight over tibia length was measured (n=10 and 8). Interventricular septum (IVS) (**D**), left ventricular posterior wall (LVPW) (**E**) thickness and left ventricular internal diameter (**F**) were measured by echocardiography in diastole. (**G**) Myh7 mRNA expression was quantified by RT-qPCR in total heart extract and normalized to 18S rRNA. (**H**) Heart cross sections were co-immunostained with anti-CX43 (in green) and anti- ACTN2 (in red) antibodies. (**I**) CX43 protein expression was analyzed by western blot and (**J**) quantified using Image J software. *: p≤0.05, ***: p≤0.001, ns: not significant (Mann Whitney test).

Also cardiomyocytes showed impaired differentiation with increased Myosin Heavy Chain 7 (Myh7) mRNA expression (Figure 4G) and decreased Connexin 43 (CX43) levels (Figure 4I-J). To test whether cardiomyocyte contractility was affected by pericyte depletion, we first measured ATPase Sarcoplasmic/Endoplasmic Reticulum Ca2+ Transporting 2 (ATP2A2) expression and found it significantly downregulated both at mRNA and protein level in Pdgfrb-DTA mice compared to control (Figure 5A-C). This was associated with diminished Phospholamban (PLN) both at Serine 16 and Threonine 17. (Figure 5D-F). On the contrary, Ryanodine Receptor 2 (RYR2) phosphorylation at serine 2814 was not significantly different in both groups (Figure 5G-H). To assess cardiomyocyte stiffness, we quantified mRNA expression of each Titin (Ttn) isoforms. Expression of stiff Ttn isoform N2B was significantly diminished while expression of Ttn N2BA isoform tended to increase. Accordingly, the N2BA/N2B ratio was significantly increased (Figure 5I-H).

**Figure 5.**
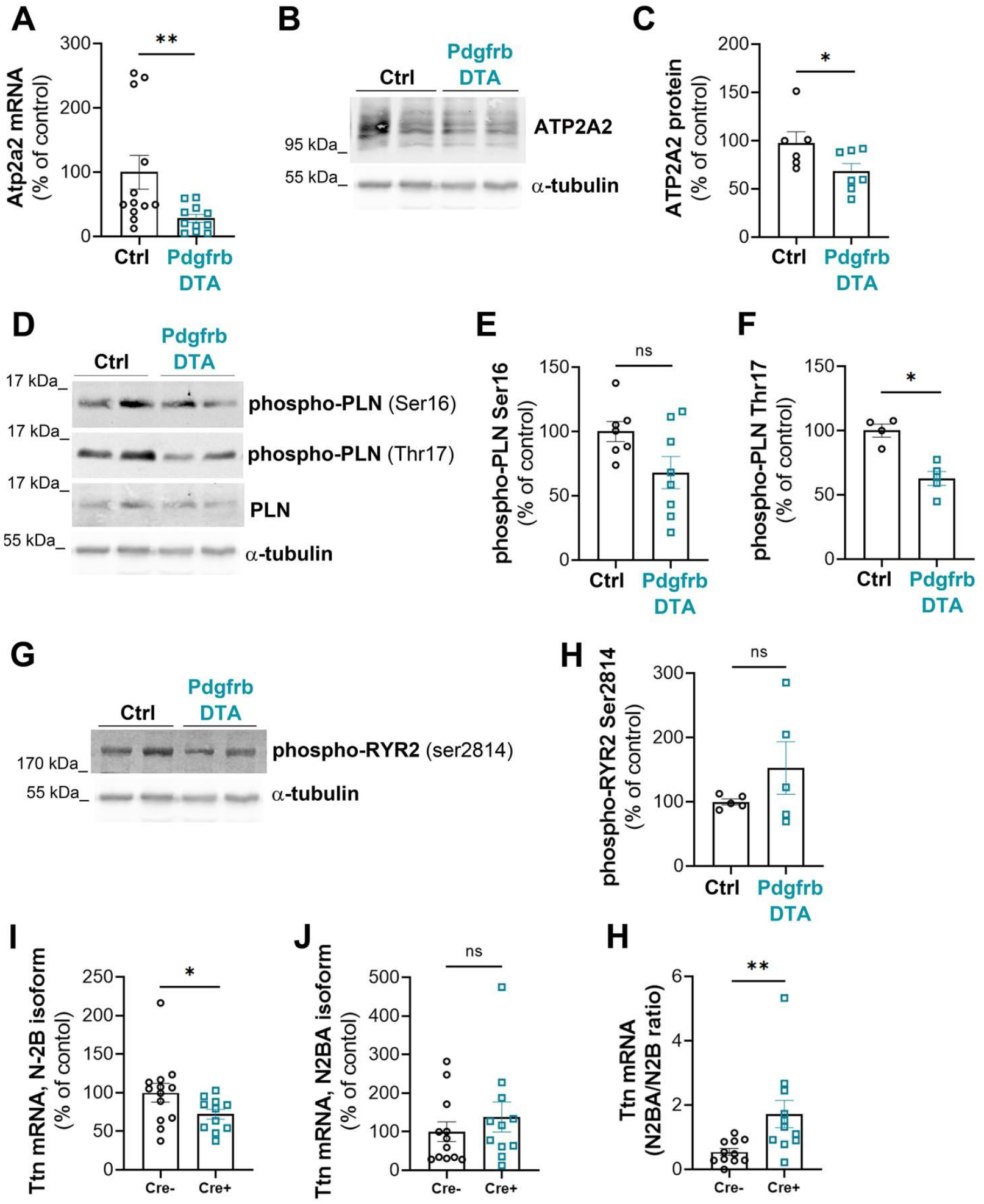
Pericyte depletion modifies Ca^2+^ homeostasis and Ttn splicing in cardiomyocytes. Pdgfrb-Cre/ERT2; RosaDTA (Pdgfrb-DTA) and RosaDTA (Control) mice were administered with tamoxifen. Mice were sacrificed 28 days after the first injection. (**A**) Atp2a2 mRNA expression was quantified by RT-qPCR in total heart extract and normalized to 18S rRNA (n=11 and 12). (**B**) ATP2A2 protein expression was analysed by western blot in total heart extract and (**C**) quantified using Image J software (n=7 and 6). (**D**) PLN phosphorylation was assessed by western blot analyses and (**E,F**) quantified using Image J software (n=5 to 8). (**G**) RYR2 phosphorylation was assessed by western blot analyses and (**H**) quantified using Image J software (n=5 in each group). Ttn, isoform N-2B (**I**) and N2BA (J) mRNA expression was quantified by RT-qPCR in total heart extract and normalized to 18S rRNA (n=11 and 12). (**H**) N2BA mRNA/N2B mRNA ratio was calculated. *: p≤0.05, **: p≤0.01, ns: not significant (Mann Whitney test).

Altogether these results demonstrate that pericyte depletion induces impaired cardiomyocyte differentiation, contractility and display a Titin isoform ratio consistent with heart failure in humans (Borbély et al. 2009). In order to test whether pericytes may affect cardiomyocyte biology directly, we performed co-culture assays. However, when co-cultured with human cardiomyocytes, mouse cardiac pericytes did modify neither ATP2A2 nor CX43 protein levels (Supplemental Figure 9) suggesting that pericytes may not affect the phenotype of cardiomyocytes directly. Impaired cardiomyocyte differentiation and contractility are then more likely the consequence of endothelium dysfunction or cardiac inflammation.

### Pericyte depletion impairs both systolic and diastolic function

Finally, in order to test whether pericyte depletion and the associated cardiac remodeling results in cardiac dysfunction, we performed echocardiography and left ventricular catheterization. As shown in figure 6, left ventricular ejection fraction (LVEF) was significantly reduced in Pdgfrb-DTA mice indicating systolic dysfunction (Figure 6A). LVEF was interestingly positively correlated with pericyte coverage (Figure 6B). Consistent with systolic dysfunction, cardiac contractility (attested by dP/dt max and contractility index) was significantly reduced (Figure 6C-E) and LV end systolic volume significantly increased (Figure 6F-G) in mice depleted from pericytes compared to control mice.

**Figure 6.**
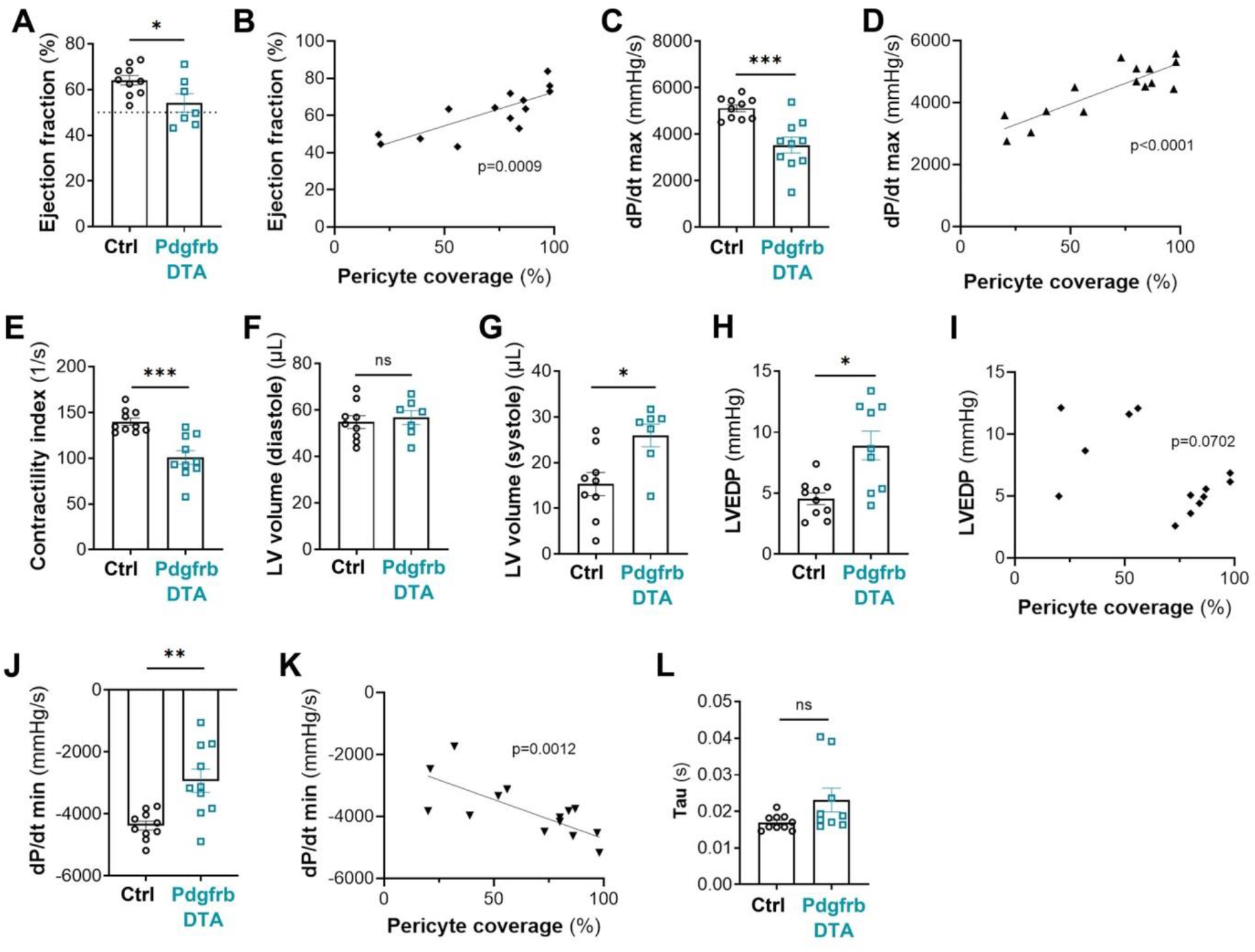
Pericyte depletion induces both systolic and diastolic dysfunction. Pdgfrb-Cre/ERT2; RosaDTA (Pdgfrb-DTA) and RosaDTA (Control) mice were administered with tamoxifen. Mice were sacrificed 28 days after the first injection. (**A**) Ejection fraction was assessed via echocardiography (n=7 and 10) and (**B**) correlated with pericyte coverage. (**C**) dP/dT max was measured by left ventricular catheterization (n=10 in each group) (**D**) correlated with pericyte coverage. **(E**) Contractility index was measured by left ventricular catheterization (n=10 in each group). LV volumes were measured in diastole (**F**) and systole (**G**) via echocardiography (n=7 and 10). (**H** LV end diastolic pressure (EDP) was measured by left ventricular catheterization (n=10 in each group) (**I**) correlated with pericyte coverage. (**J**) dP/dT min was measured by left ventricular catheterization (n=10 in each group) (**K**) correlated with pericyte coverage. (**L**) Tau was measured by left ventricular catheterization (n=10 in each group). *: p≤0.05, **: p≤0.01, ***: p≤0.001, ns: not significant (Mann Whitney test).

Mice depleted from cardiac pericytes also displayed diastolic dysfunction attested by increased end diastolic pressure (EDP) (Figure 6H,I) and decreased dP/dt min (Figure 6J-K). Relaxation constant Tau tended to increase in Pdgfrb-DTA mice but did not reach significance (Figure 6L).

In conclusion, the present study demonstrates for the first time that pericytes are necessary for proper cardiac function.

## Discussion

The present study demonstrates for the first time that, just like in the brain, pericytes are necessary for endothelium integrity and immune quiescence in the heart. Importantly, in this organ, pericytes are not only necessary for coronary vasculature integrity but also for proper cardiac function suggesting that modifications of pericyte properties may participate in the pathophysiology of cardiac diseases.

Besides, this study reveals that cardiac pericytes may undergo strong remodeling upon injury (e.i. renewal, expression of SMA). This is actually consistent with what has been observed in obese and hypertensive ZSF1rats (van Dijk et al. 2016) in which pericytes were shown to be disorganized and more numerous. This piece of data actually suggests that, even though pericytes may induce similar effects in the brain and in the heart by maintaining microvasculature integrity, pericyte turnover and biology in these organs may be very different. Indeed, in the central nervous system, pericyte impairment seems to be essentially characterized by pericyte loss. Notably, one of the earliest hallmarks of diabetic retinopathy is the loss of pericytes. The diabetic microenvironment is suggested to be particularly detrimental to pericyte survival in the retina since during the early stages of retinopathy, the pericyte to EC ratio decreases from 1:1 to 1:4 (Robison, Kador, and Kinoshita 1985). In the brain, pericyte number and coverage in the cortex and hippocampus of subjects with Alzheimer’s disease compared with neurologically intact controls were shown to be reduced by 59% and 60% (Sengillo et al. 2013). Similarly, in the spinal cord, immunostaining for PDGFRβ, have indicated a 54 % (p < 0.01) reduction in pericyte number in amyotrophic lateral sclerosis patients compared to controls (Winkler et al. 2013). However, in the heart, pericyte impairment is more likely characterized by phenotypic changes with acquisition of an ‘activated’ phenotype. Notably, the number of pericytes was shown to be increased in the heart of mice infused with angiotensin-2.

Moreover these pericytes express higher levels of SMA (Su et al. 2020). As mentioned above, cardiac pericytes were shown to be disorganized and more numerous in obese and hypertensive ZSF1 rats (van Dijk et al. 2016). In our model, pericyte depletion induced pericyte activation and renewal.

Pericyte loss have been previously associated with reduced LVEF (Chintalgattu et al. 2013) (He, Zeng, and Chen 2016), dilated cardiomyopathy, cardiac fibrosis, contractile dysfunction (Siao et al. 2012) and cardiac hypertrophy (Tao et al. 2017). Our study confirms that pericytes are necessary for proper cardiac contractility and systolic function, while pericyte loss does not seem to promote cardiac fibrosis, hypertrophy or dilation. In Siao et al. study, cardiac fibrosis and dilation may then be due to the effect of the mutant form of ProNGF on other cell types, notably cardiomyocytes and/or fibroblasts (Siao et al. 2012), while in Tao et al. study (Tao et al. 2017), cardiac hypertrophy may be due to Notch3 deletion in cardiomyocytes (Øie et al. 2010).

Whether or not pericytes may dialogue with cardiomyocytes directly remains to be further investigated, while our data suggest that pericyte associated deregulation of ATP2A2 and CX43 expression in cardiomyocyte require at least a third cell type, another study suggest that pericytes may dialogue with cardiomyocytes directly via MiR-132 transfer (Katare et al. 2011).

One of the main limitations of this study is technic because a mouse model only targeting cardiac pericytes still needs to be generated to obtain robust and specific characterization of the role of cardiac pericytes. Indeed, Pdgfrb promoter does not only target pericytes but also SMCs and Pdgfrb promoter does not only target cardiac mural cells but mural cells of the entire body. However, several mouse models targeting pericytes have been used especially in the brain and most of these models are based on PDGFB-PDGFRB signaling which is crucial for pericyte recruitment and survival (Hoch and Soriano 2003). At first, developmental studies were performed on *Pdgfb* (Levéen et al. 1994) and *Pdgfrb* (Soriano 1994) deficient embryos, which are not viable. Then mice harboring mutations within the *Pdgfrb* gene (the *Pdgfrb*^redeye/redeye^ mutant (Jadeja et al. 2013) and the *Pdgfrb*^F7/F7^ mutant (Angeliki Maria Nikolakopoulou et al. 2017)) have been used. These mutants display progressive pericyte loss especially in the central nervous system (brain and retina). Mouse models with inducible pericyte depletion have been developed, these models are tamoxifen inducible, and use either Diphtheria toxin A (DTA) or its receptor DTR to induce cell ablation (Eilken et al. 2017, 1). These last models allow the exploration of the role of pericytes in adults excluding the potential consequences of developmental defects. The Pdgfrb-Cre/ERT2; Rosa iDTR model is proposed to induce a less severe pericyte depletion than the Pdgfrb-Cre/ERT2; Rosa iDTA model however, it requires administration of both tamoxifen and Diphtheria toxin (Eilken et al. 2017, 1). Alternatively to the *Pdgfrb* promoter, the *Cspg4* (also known as NG2) promoter has been proposed to induce pericyte depletion. However, its induction efficiency is lower than the one of Pdgfrb and just like Pdgfrb promoter, it induces recombination in SMCs (Mayr et al. 2022, 2). Finally, a strategy utilizing a double-promoter approach with Pdgfrb and the Cspg4 have been used (Angeliki M. Nikolakopoulou et al. 2019) to target pericytes more specifically, however both of these promoters actually also target SMCs as described above.

## Acknowledgments

We thank Annabel Reynaud, Sylvain Grolleau, and Maxime David for their technical help. We thank Christelle Boullé for administrative assistance.

