## Supplemental data for "Cardiac pericytes are necessary for coronary vasculature integrity and cardiomyocyte differentiation"

### Supplementary data

**Supplemental tables**

A/ List of primary antibodies

| Target antigen | Vendor or Source | Catalog # | Working concentration |
|---|---|---|---|
| mouse PECAM-1 (CD31) | BMA Biomedicals | T-2001 | 2 µg/mL |
| mouse PODXL | R&D systems | AF1556 | 4 µg/mL |
| mouse Cy3-anti-smα-actin | Sigma | C6198 | 5 µg/mL |
| mouse NG2 | Chemicon | AB 5320 | |
| mouse ACTN2 | Sigma | A 2172 | |
| mouse VIM | Cell signaling technology | 5741 | |
| mouse ICAM-1 (CD54) | BD Pharmingen | 550287 | 2.5 µg/mL |
| mouse VCAM-1 | Invitrogen | 14-1061-82 | |
| mouse CDH5 | R&D systems | AF1002 | |
| mouse FGB | Abcam | ab227063 | |
| mouse COL1A1 | Abcam | ab21285 | |
| mouse CX43 | Sigma | C6219 | |
| mouse CD41 | Proteintech | 18308-1-AP | 3.5 µg/mL |
| mouse Myh7 | Atlas antibodies | HPA001239 | 1 µg/mL |
| mouse Desmin | DB Biotech | DB148-0.1 | 0.5 µg/mL |
| mouse CD45 | BD Pharmingen | 550539 | 0.625 µg/mL |
| mouse CD68 | Biolegend | 137001 | 5 µg/mL |
| mouse CD3 | Santa-Cruz Biotechnology | sc-1127 | |
| mouse B220 | R&D systems | MAB1217 | |
| GFP | Novus | NB100-1770SS | |
| GFP | Invitrogen | A6455 | |
| mouse ATP2A2 | Badrilla | A010-80 | |
| phospho-PLN (Ser16) | Badrilla | A010-12 | |
| phospho-PLN (Thr17) | Badrilla | A010-13 | |
| total-PLN | Badrilla | A010-14 | |
| phospho-RYR2 (Ser2814) | Badrilla | A010-31 | |
| α-tubulin | Sigma | T5168 | |
| human CDH5 | Cell signaling technology | 2500 | |
| human ICAM-1 | Santa-Cruz Biotechnology | sc-8439 | |

B/ List of secondary antibodies

| Target antigen | Conjugate | Vendor or Source | Catalog # | Working concentration |
|---|---|---|---|---|
| Rat IgG | Biotin | Jackson ImmunoResearch | 712-065-153 | Dilution 1/500 |
| Rabbit | Biotin | Jackson ImmunoResearch | 711-065-152 | Dilution 1/500 |
| Goat IgG | Alexa Fluor 568 | Invitrogen | A-11057 | 10 µg/mL |
| Rat IgG | Alexa Fluor 647 | Invitrogen | A-48265 | 10 µg/mL |
| Hamster IgG | Biotin | Jackson ImmunoResearch | 127-065-160 | Dilution 1/500 |
| Goat IgG | Alexa Fluor 488 | Invitrogen | A-11055 | 10 µg/mL |
| Rabbit IgG | Alexa Fluor 488 | Invitrogen | A-21206 | 10 µg/mL |
| Rabbit IgG | Alexa Fluor 568 | Invitrogen | A-10042 | 10 µg/mL |

**Supplemental Table 1: List of antibodies used for immunostainings**

| | | |
|---|---|---|
| mouse 18S | F | 5′-CGCGGTTCTATTTTGTTGGT-3′ |
| | R | 5′-AGTCGGCATCGTTTATGGTC-3′ |
| mouse Col1a1 | F | 5′-CAACCTCAAGAAGGCCCTGC-3′ |
| | R | 5′-TGTCCAAGGGAGCCACATCG-3′ |
| mouse Col3a1 | F | 5′-AGCACGAGGTCTTGCTGGAC-3′ |
| | R | 5′-ACCAGCTGTACCAGGCTGAC-3′ |
| mouse Tgfb1 | F | 5′-GCTAATGGTGGACCGCAACAAC-3′ |
| | R | 5′-CACTGCTTCCCGAATGTCTGAC-3′ |
| mouse Myh7 | F | 5′-GGATGACGTCACCTCCAACA-3′ |
| | R | 5′-AGATCAGAGCCTCCTTCTCGT-3′ |
| mouse Ctgf | F | 5′-GACCCAACTATGATGCGAGCC-3′ |
| | R | 5′-TCCCACAGGTCTTAGAACAGG-3′ |
| mouse Ttn N-2B | F | 5′-ACAGTGGGAAAGCAAAGACATC-3′ |
| | R | 5′-AGGTGGCCCAGAGCTACTTC-3′ |
| mouse Ttn N2BA | F | 5′-GAGACATTGCTCCGCTTTTC-3′ |
| | R | 5′-GATCTCCAAAGAGGCTGTC-3′ |
| mouse Atp2a2 (Serca2a) | F | 5′-GATCCTCTACGTGGAACCTTTG-3′ |
| | R | 5′-GGTAGATGTGTTGCTAACAACG-3′ |
| human ACTB (β-actin) | F | 5′-GGAGGAGCTGGAAGCAGCC-3′ |
| | R | 5′-GCTGTGCTACGTCGCCCTG-3′ |
| human ICAM-1 | F | 5′-ACGCCGGAGGACAGGGCATT-3′ |
| | R | 5′-GGGGCTATGTCTCCCCCACCA-3′ |
| human VCAM-1 | F | 5′-GGCCCAGTTGAAGGATGCGGG-3′ |
| | R | 5′-AGAGCACGAGAAGCTCAGGAGAA-3′ |
| human CDH2 | F | 5′-CCGGTTTCATTTGAGGGCAC-3′ |
| | R | 5′-CCCATTGAGGGCATTGGGAT-3′ |
| human IL-6 | F | 5′-GAAGATTCCAAAGATGTAGCCGC-3′ |
| | R | 5′- GGTTGTTTTCTGCCAGTGCCTC-3′ |

**Supplemental Table 2: List of primers used for reverse transcription (RT) quantitative polymer chain reaction (qPCR).** F: forward; R: reverse

**Supplemental figures and figure legends**

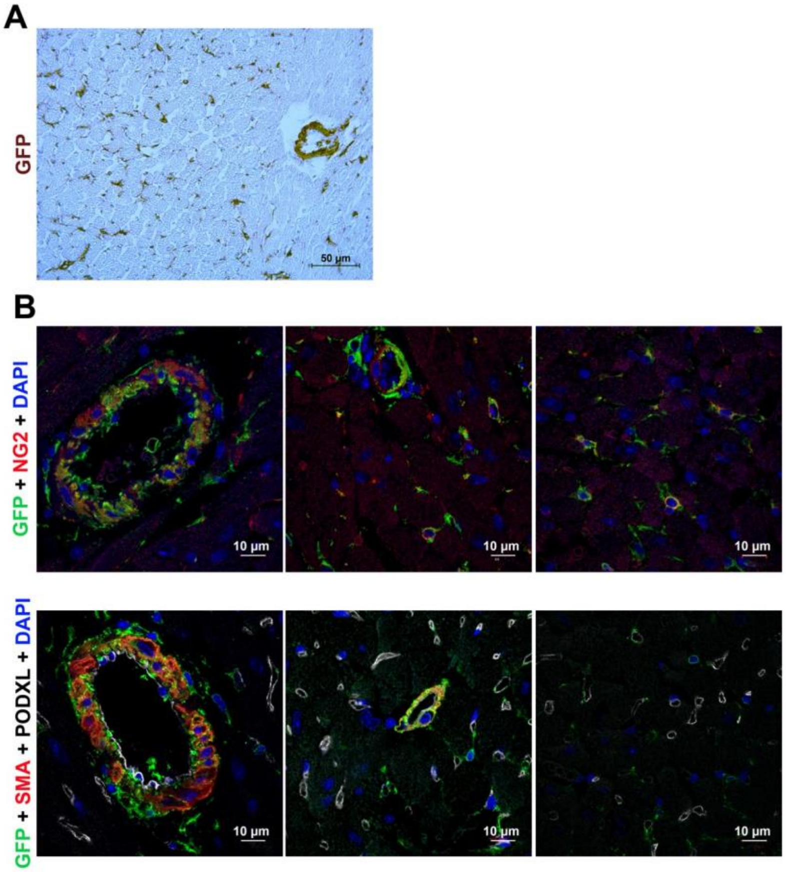

**Supplemental Figure 1:** (**A-B**)**:** Pdgfrb-Cre/ERT2; Rosa-mTmG mice were administered with tamoxifen and sacrificed 7 days after the first injection. (**A**) Heart cross sections were immunostained with anti-GFP antibodies (in brown). (**B**) Heart cross sections were co-immunostained with anti-GFP antibodies (in green) together with anti-NG2 antibodies (in red) to identify pericytes or anti-SMA (in red) and anti-PODXL antibodies (in white) to identify SMCs and endothelial cells respectively.

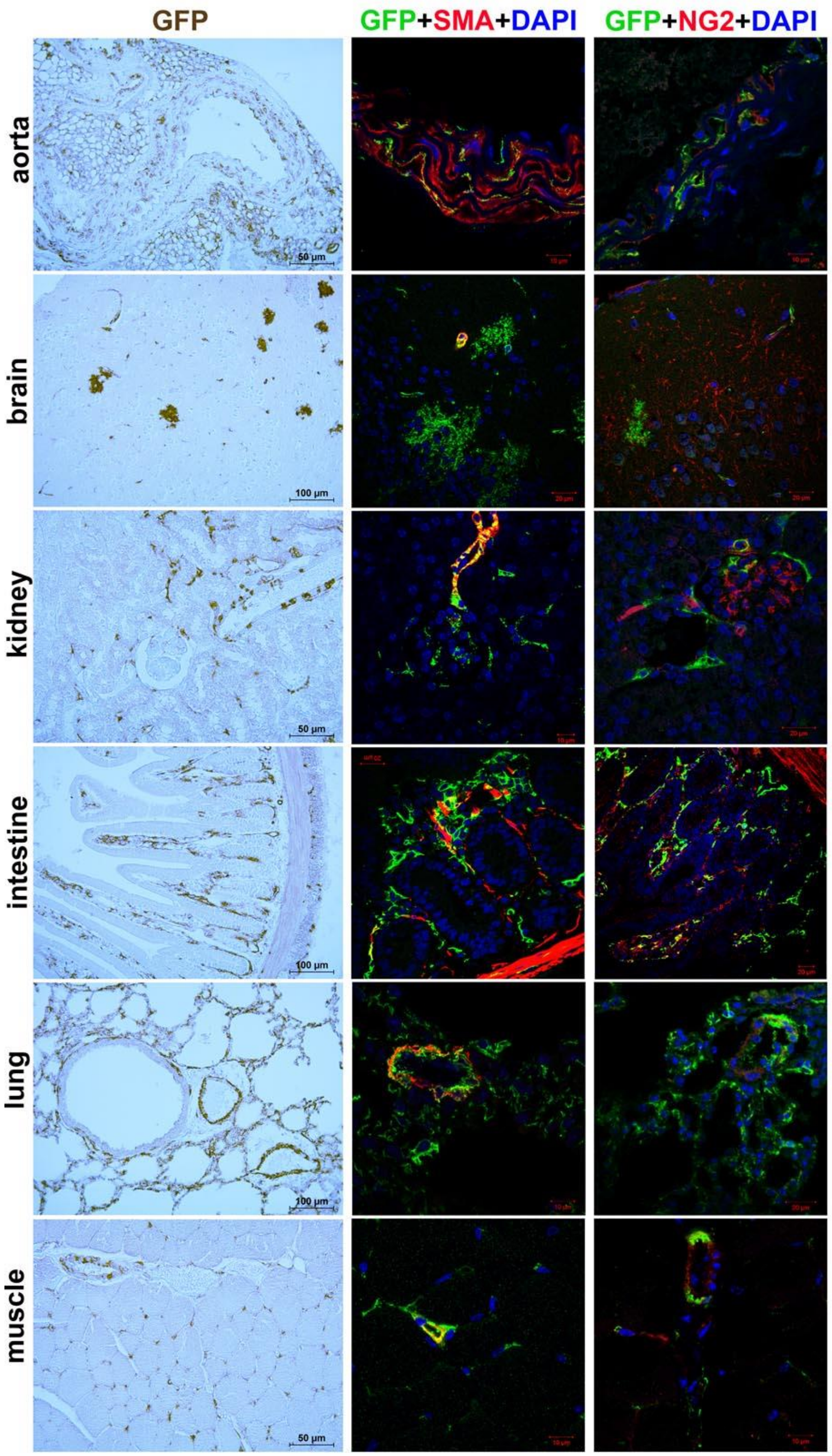

**Supplemental Figure 2:** Pdgfrb-Cre/ERT2; Rosa-mTmG mice were administered with tamoxifen and sacrificed 7 days after the first injection. (**A**) Tissue sections were immunostained with anti-GFP antibodies alone or together with anti-SMA antibodies to identify SMCs or anti-NG2 antibodies to identify pericytes.

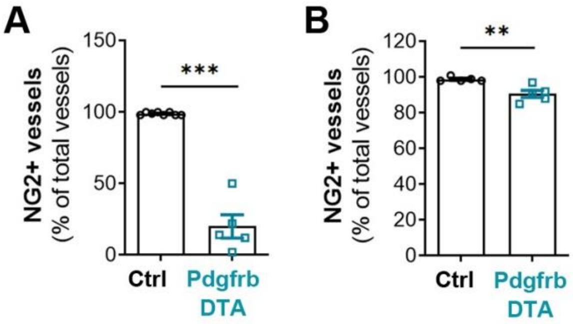

**Supplemental Figure 3:** Pdgfrb-Cre/ERT2; RosaDTA (Pdgfrb-DTA) mice were administered with tamoxifen. Mice were sacrificed 7 days (**A**) or 28 days after the first injection (**B**). Heart cross sections were co-immunostained with anti-NG2 antibodies (in red) together with anti-PODXL antibodies (in green), the percentage of NG2 positive vessel was counted. **: $p \leq 0.01$, ***: $p \leq 0.001$ (Mann Whitney test).

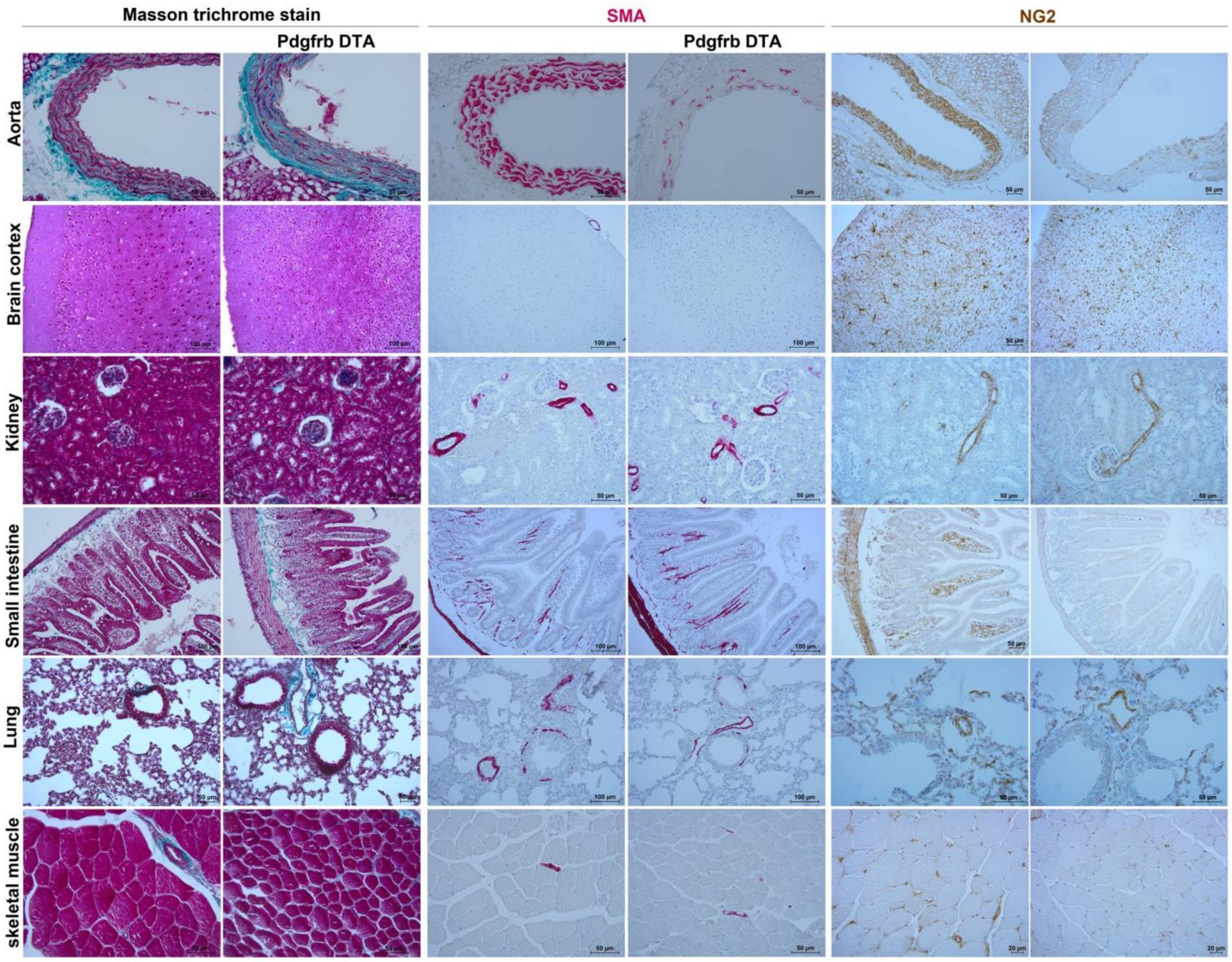

**Supplemental Figure 4:** Pdgfrb-Cre/ERT2; RosaDTA (Pdgfrb-DTA)mice were administered with tamoxifen and sacrificed 28 days after the first injection. Tissues sections were staining with Masson's trichrome stain, anti-SMA or anti-NG2 antibodies.

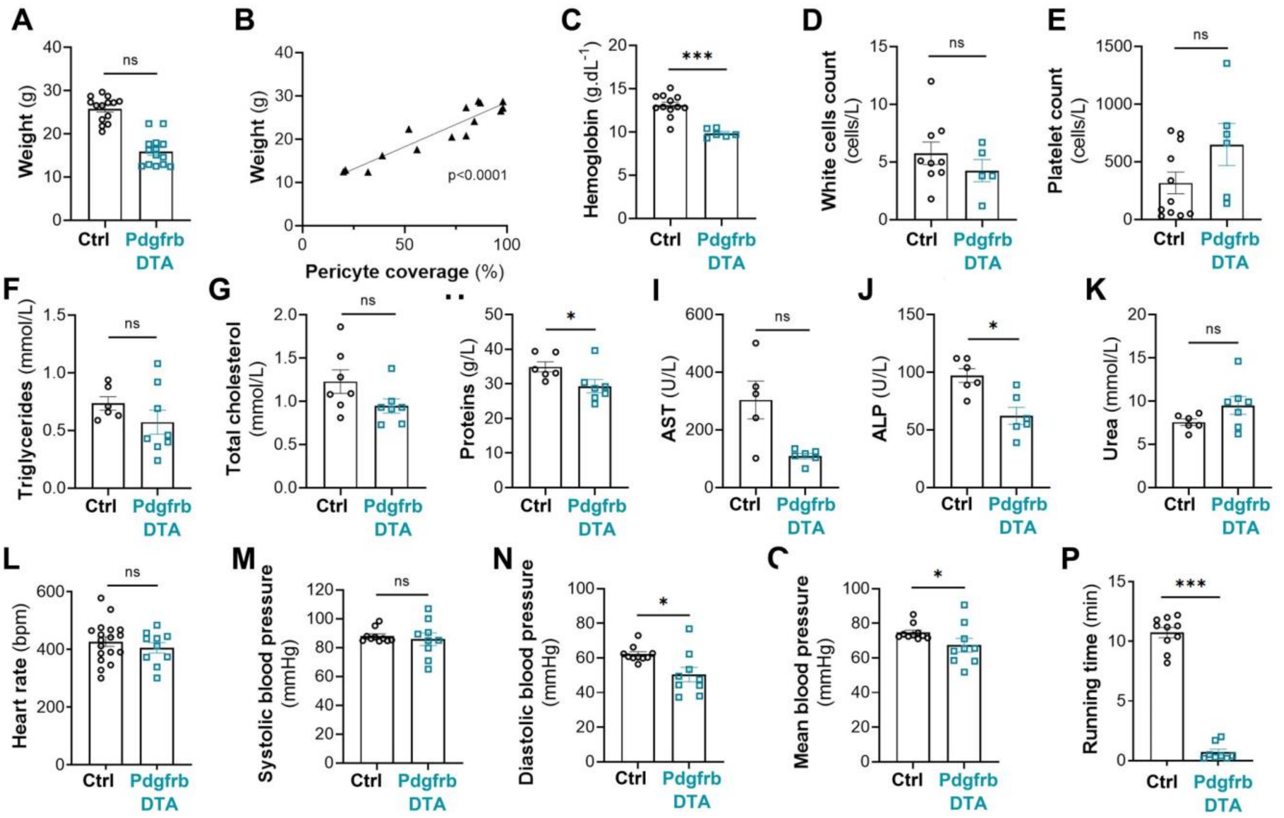

**Supplemental Figure 5:** Pdgfrb-Cre/ERT2; RosaDTA (Pdgfrb-DTA) and RosaDTA (Control) mice were administered with tamoxifen. Mice were sacrificed 28 days after the first injection. (A) The weight of mice was measured and (**B**) correlated with pericyte coverage. (**C**) Hemoglobin content, (**D**) white blood cells count and platelet count (**E**) was calculated in total blood samples. (**F**) Triglycerides, (**G**) total cholesterol, (**H**) proteins, (**I**) Aspartate aminotransferase (AST), (**J**) Alcaline phosphatase (ALP) and (**K**) urea levels were measured in plasma samples. (**L**) Heart rate, (M) systolic, (N) diastolic and (**O**) mean blood pressures were measured via left ventricular catheterization. (P) Exercise tolerance on a treadmill was assessed. *: p≤0.05, ***: p≤0.001, ns: not significant (Mann Whitney test).

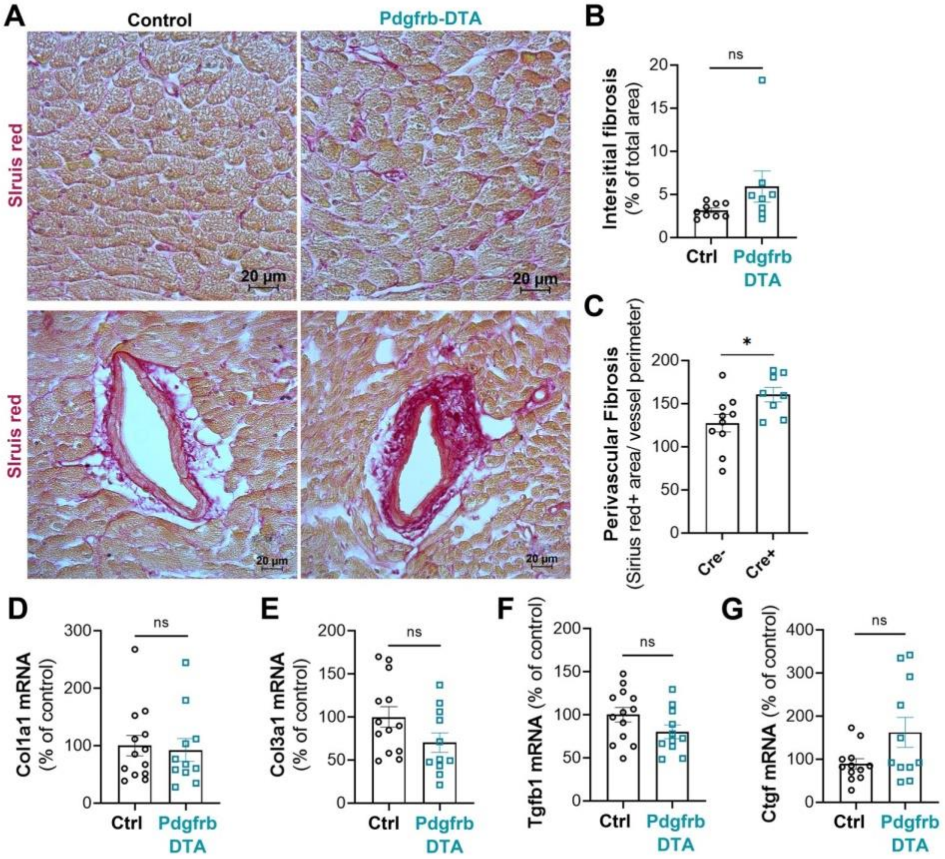

**Supplemental Figure 6:** Pdgfrb-Cre/ERT2; RosaDTA (Pdgfrb-DTA) and RosaDTA (Control) mice were administered with tamoxifen. Mice were sacrificed 28 days after the first injection. (**A**) Heart cross sections were stained with Sirius red to identify fibrosis. (**B**) Interstitial fibrosis was quantified using Image J software (n=9 and 11). (**C**) Perivascular fibrosis was quantified using Image J software (n=9 and 11). Col1a1 (D), Col3a1 (E), Tgfb1 (F) and Ctgf (G) mRNA was quantified by RT-qPCR in total heart extracts and normalized to 18S rRNA. *: p≤0.05, ns: not significant (Mann Whitney test).

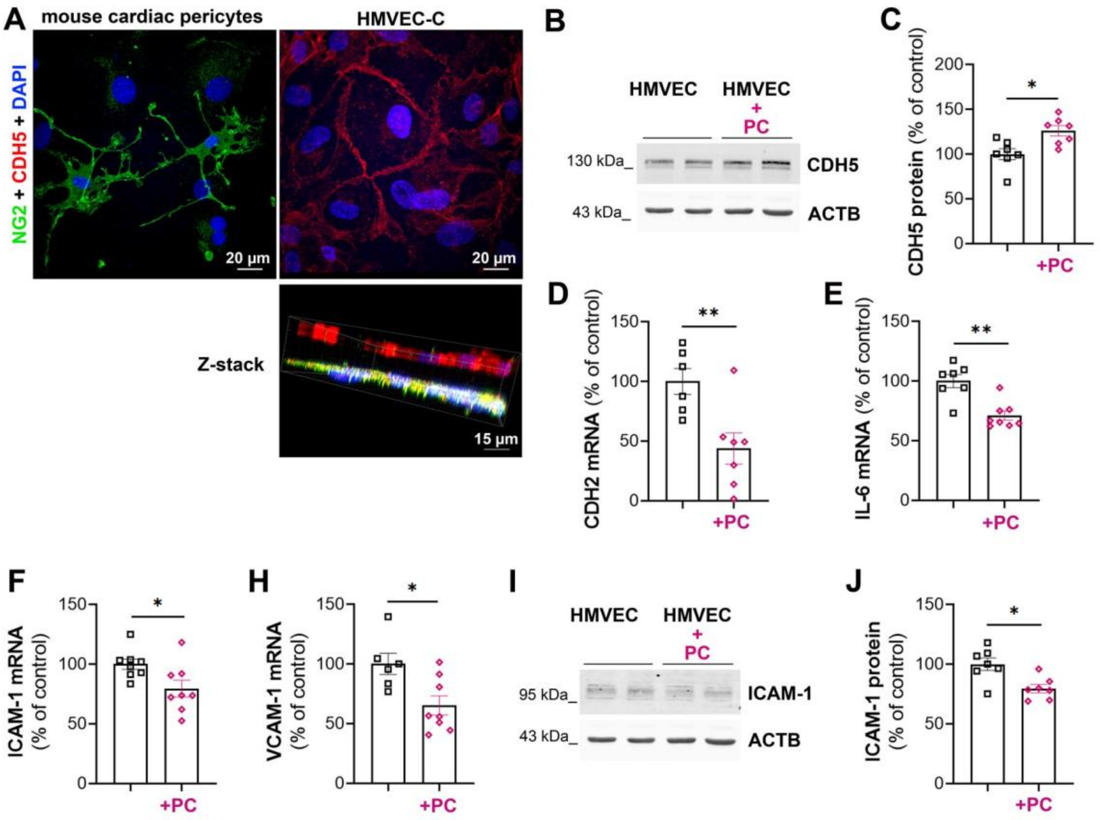

**Supplemental Figure 7:** Cardiac HMVECs were co-cultured or not with mouse cardiac pericyte. Each cell type was plated on each side of a 0.4 µm pored Transwell® membrane for 48 hours. (**A**) Pericytes were immunostained with anti-NG2 antibodies in green, HMVECs were immunostained with anti-CDH5 antibodies in red. (**B**) CDH5 protein level in HMVECs was evaluated via western blot analysis and (**C**) quantified using Image J software. *CDH2* (**D**), *IL-6* (**E**), *ICAM-1* (**F**) and *VCAM-1* (**H**) mRNA levels were quantified via RT-qPCR and normalized to *ACTB* mRNA. (**I**) ICAM-1 protein level in HMVECs was evaluated via western blot analysis and (**J**) quantified using Image J software. *: p≤0.05, **: p≤0.01 (Mann Whitney test).

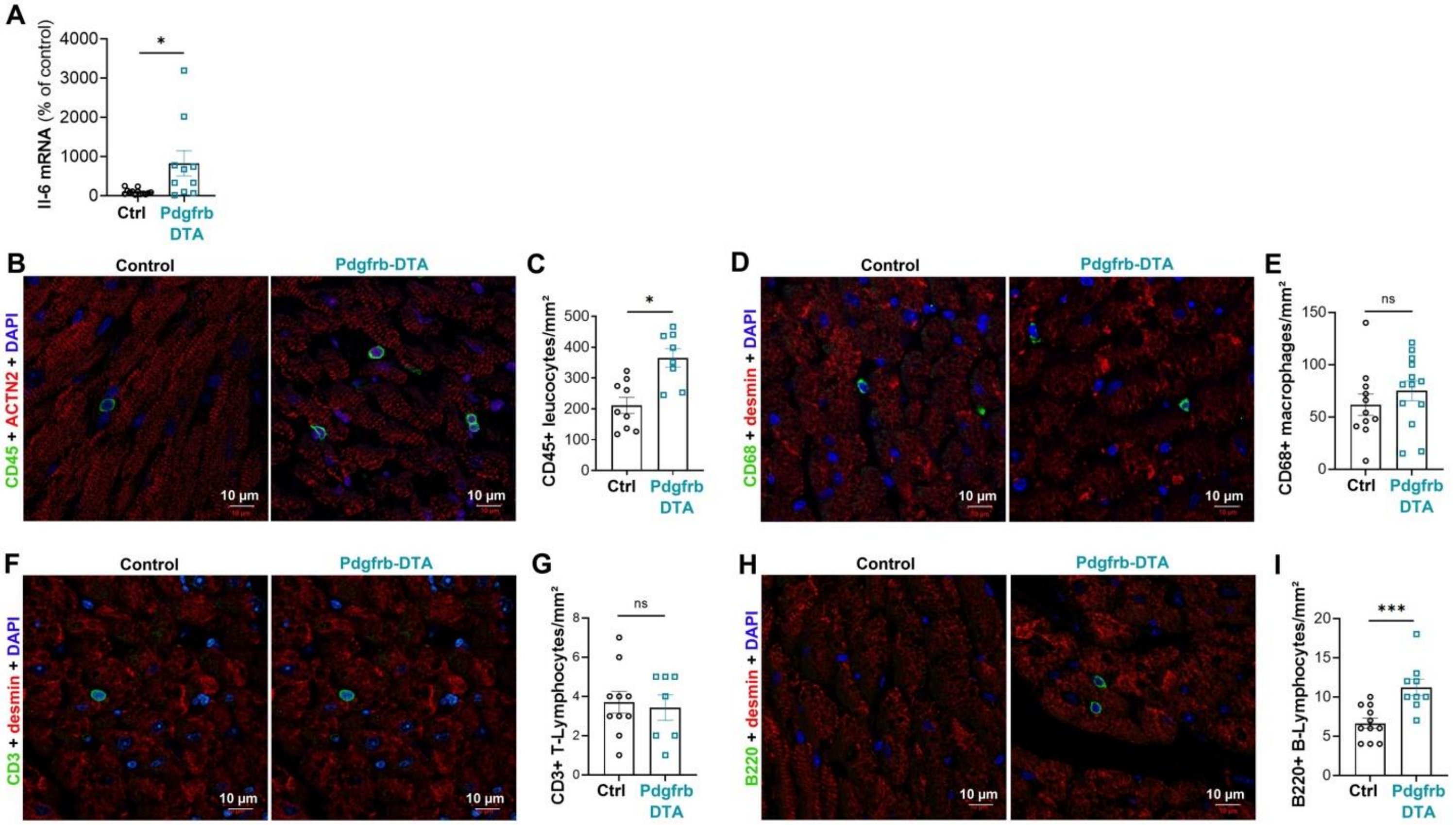

**Supplemental Figure 8:** Pdgfrb-Cre/ERT2; RosaDTA (Pdgfrb-DTA) and RosaDTA (Control) mice were administered with tamoxifen. Mice were sacrificed 28 days after the first injection. (**A**) *Il-6* mRNA level was quantified via RT-qPCR and normalized to 18S rRNA. (**B**) Heart cross sections were co-immunostained with anti-CD45 antibodies (in green) and anti-ACTN2 antibodies (in red). (**C**) The number of CD45 positive leucocytes/$mm^2$ was quantified using Image J software. (**D**) Heart cross sections were co-immunostained with anti-CD68 antibodies (in green) and anti-desmin antibodies (in red). (**E**) The number of CD68 positive macrophages/$mm^2$ was quantified using Image J software. (**F**) Heart cross sections were co-immunostained with anti-CD3 antibodies (in green) and anti-desmin antibodies (in red). (**G**) The number of CD3 positive T-lymphocytes/$mm^2$ was quantified using Image J software. (**H**) Heart cross sections were co-immunostained with anti-B220 antibodies (in green) and anti-desmin antibodies (in red). (**I**) The number of B220 positive B-lymphocytes/$mm^2$ was quantified using Image J software. *: $p \leq 0.05$, ***: $p \leq 0.001$, ns: not significant (Mann Whitney test).

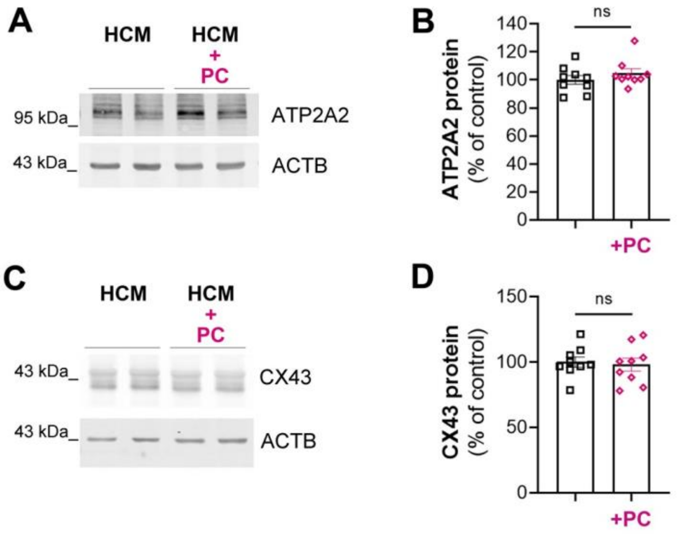

**Supplemental Figure 9:** Human cardiac myocyte (HCM) were co-cultured or not with mouse cardiac pericytes. Each cell type was plated on each side of a 0.4 µm pored Transwell® membrane for 48 hours. (**A**) ATP2A2 protein level in HCMs was evaluated via western blot analysis and (**B**) quantified using Image J software. (**C**) CX43 protein level in HCMs was evaluated via western blot analysis and (**J**) quantified using Image J software. ns: not significant (Mann Whitney test).
